# Integration of plasticity mechanisms within a single sensory neuron of *C. elegans* actuates a memory

**DOI:** 10.1101/157958

**Authors:** Josh D. Hawk, Ana C. Calvo, Agustin Almoril-Porras, Ahmad Aljobeh, Maria Luisa Torruella-Suárez, Ivy Ren, Nathan Cook, Joel Greenwood, Linjiao Luo, Aravinthan D.T. Samuel, Daniel A. Colón-Ramos

**Author notes:** To whom correspondence should be addressed: Daniel A. Colón-Ramos, Ph.D. Department of Cell Biology Department of Neuroscience Yale Program in Cellular Neuroscience, Neurodegeneration and Repair Yale University School of Medicine 333 Cedar St SHM B163D New Haven, CT 06510.

## Abstract

Neural plasticity—the ability of a neuron to change its cellular properties in response to past experiences—underpins the nervous system’s capacity to form memories and actuate behaviors. How different plasticity mechanisms act together *in vivo* and at a cellular level to transform sensory information into behavior is not well understood. Here we show that in the nematode *C. elegans* two plasticity mechanisms—sensory adaptation and presynaptic plasticity—act within a single cell to encode thermosensory information and actuate a temperature-preference memory. Sensory adaptation enables the primary thermosensory neuron, AFD, to adjust the temperature range of its sensitivity to the local environment, thereby optimizing its ability to detect temperature fluctuations associated with migration. Presynaptic plasticity transforms this thermosensory information into a behavioral preference by gating synaptic communication between sensory neuron AFD and its postsynaptic partner, AIY. The gating of synaptic communication is regulated at AFD presynaptic sites by the conserved kinase nPKCε. Bypassing or altering AFD presynaptic plasticity predictably changes the learned behavioral preferences without affecting sensory responses. Our findings indicate that two distinct and modular neuroplasticity mechanisms function together within a single sensory neuron to encode multiple components of information required to enact thermotactic behavior. The integration of these plasticity mechanisms result in a single-cell logic system that can both represent sensory stimuli and guide memory-based behavioral preference.

## Introduction

When an animal experiences a favorable condition, it can form a memory that guides its future behavior. The future performance of these learned behavioral preferences requires the animal to navigate sensory-rich environments, extract behaviorally-relevant information and differentially act based on its previous experience. While it is widely accepted that plasticity mechanisms underpin experience-driven behaviors, how different plasticity mechanisms act *in vivo* and within single cells to facilitate recall and actuation of learned behavioral preferences is not well understood (Basu and Siegelbaum, 2015; Davis, 2011; Gjorgjieva et al., 2016; Mayford et al., 2012).

The nematode *C. elegans* does not have an innate preferred temperature. Instead, *C. elegans* cultivated at a given temperature will remember this temperature (T_c_) and migrate towards it when on a thermal gradient (Hedgecock and Russell, 1975). This memory can be trained to a new temperature within four hours (Biron et al., 2006; Chi et al., 2007; Hedgecock and Russell, 1975; Mohri et al., 2005; Ramot et al., 2008b) (Figure 1B-D). Neurons specifically required for thermotaxis behavior have been identified by neuronal ablation studies (Beverly et al., 2011; Biron et al., 2008; Kuhara et al., 2008; Mori and Ohshima, 1995), and their connectivity is known (White et al., 1986). From this work, we understand that temperature preference depends on a thermosensory neuron, called AFD, which has specialized molecular pathways that allow it to respond to temperature changes as small as 0.1°C (Clark et al., 2006; Kimura et al., 2004; Mori and Ohshima, 1995; Ramot et al., 2008a). AFD responses are observed only near an adaptable sensory threshold that correlates, on long timescales, with the cultivation temperature memory. These observations led to the hypothesis that the AFD thermosensory threshold represents the memory for the preferred temperature (Aoki and Mori, 2015; Biron et al., 2006; Garrity et al., 2010; Kimura et al., 2004; Luo et al., 2014; Yu et al., 2014). AFD has also been observed to adapt its thermosensory threshold on short timescales (Ramot et al., 2008a; Yu et al., 2014), but the behavioral repercussions of this more rapid adaptation have not been examined. Thus, the role of the adaptable AFD sensory threshold in the actuation of the cultivation temperature memory is not well understood.

**Figure 1.**
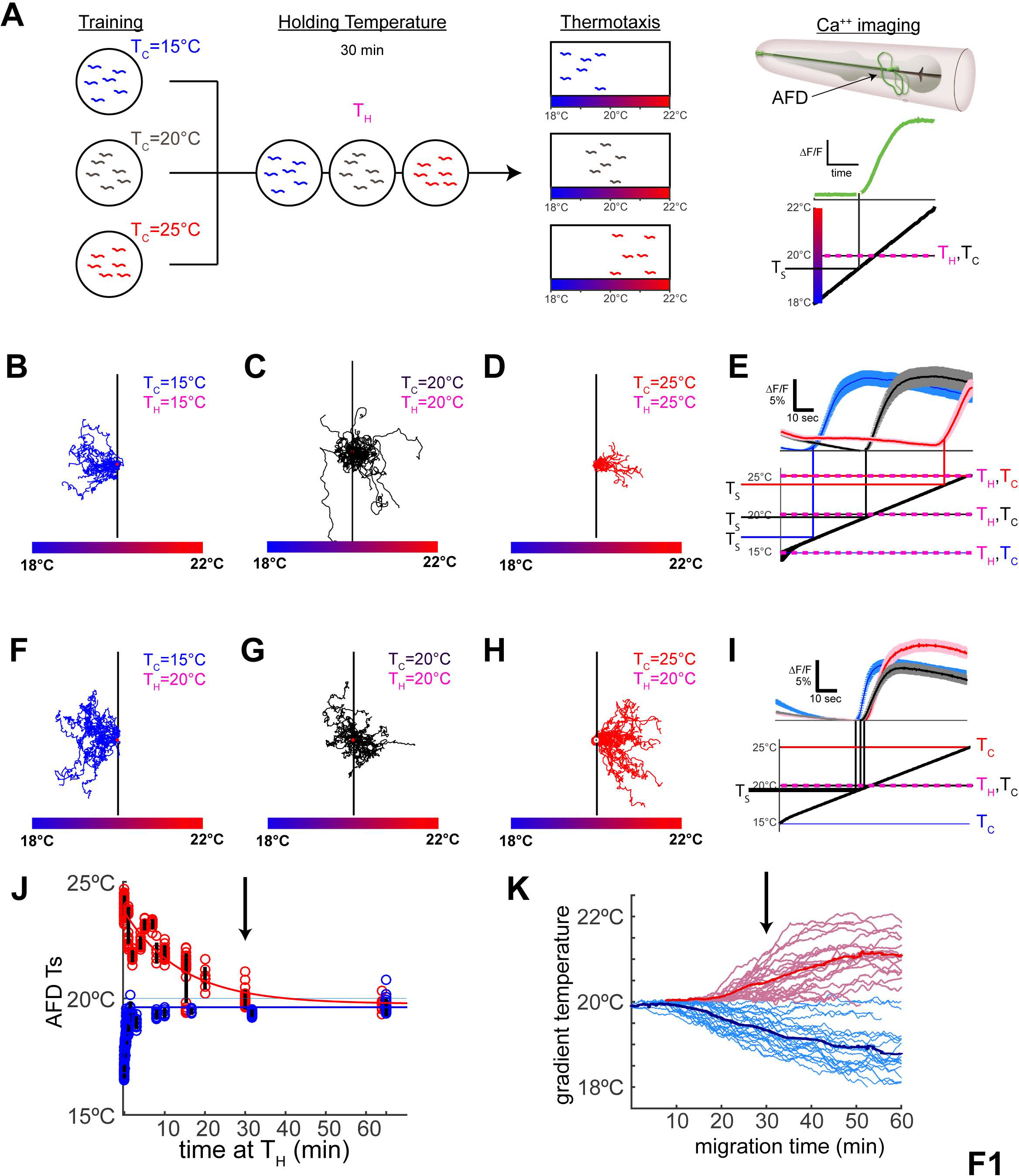
AFD thermosensory threshold adapts within minutes and does not represent the temperature preference memory. (**A**) Schematic illustration of the experimental paradigms. Animals cultivated for 4hr either at 15°C, 20°C or 25°C (“Training”) were then held at specified temperatures (“Holding Temperature”, T_H_) for 30 minutes prior to being tested in behavioral assays (“Thermotaxis”) or calcium imaging in AFD (“Ca^++^ imaging”). The response of a single AFD neuron (green trace) is illustrated as a change in fluorescence (∆F/F) over time when applying a linear thermal ramp (thick black diagonal line) from 18°C to 22°C. (**B-E**) Worms trained at 15°C, 20°C, or 25°C were examined for thermotaxis behavior in **B-D** and calcium imaging in AFD in **E**. The mean AFD response threshold for T_C_=15°C was 16.9°C +/− 0.5°C (blue trace), T_C_=20°C was 19.9°C +/− 0.5°C (black trace), and T_C_=25°C was 23.9°C +/− 0.4°C (red trace). Note that when T_H_=T_C_, AFD response threshold (T_S_) correlates with the T_C_, consistent with previous reports (Clark et al., 2006; Kimura et al., 2004; Ramot et al., 2008a). Traces for individual animals in Figure S1B-C. (**F-I**) As in **B-E**, but animals were held at T_H_=20°C for 30 minutes prior to thermotaxis behavior in **F-H** or calcium imaging in **I**. Note that, regardless of T_H_, animals retain the cultivation temperature memory and perform the thermotaxis behavior (**F-H**) as in **B-D**. (**I**) In contrast, the AFD response threshold (T_S_) adapts to the holding temperature (T_H_) and all animals, regardless of cultivation temperature memory, respond near T_H_=20°C: T_S_=19.4°C +/−0.1°C for T_C_=15°C, 19.8°C +/− 0.2°C for T_C_=20°C, and 19.9°C +/−0.3°C for T_C_=25°C. Traces for individual animals are in Figure S1E-G. (**J**) AFD adaption kinetics were assessed in a training paradigm as described in **A**, but with varying amounts of time held at T_H_. The threshold T_S_ is plotted for each duration of holding at T_H_. The time course of adaptation in AFD for cooling temperatures (T_H_=20°C & T_C_=25°C, tau=14min) is slower than that observed for warming temperatures (T_H_=20°C & T_C_=15°C, tau=45sec). Yet, note that by 30 minutes (black arrow) sensory threshold responses in AFD have adapted to very near T_H_ regardless of T_C_. See also Figure S1H-D’. (**K**) Behavioral kinematics showing migration over time in animals illustrate that migration onset correlates well with AFD adaptation kinetics, with peak migration occurring around 30 min after worms were placed at 20°C (black vertical arrow). See also Figure S1E’. Error bars denote SEM.

## Results

To better understand AFD sensory responses in relationship to thermotaxis behavior, we developed a custom thermoelectric control system (see Materials and Methods and Figure S1A) that can rapidly (~1°C/second) and precisely (+/− 0.02°C) control temperature while we image calcium dynamics using GCaMP6 (Chen et al., 2013) in individual neurons of live animals. Implementing similar conditions to those used in published studies, we reproduced the observation that sensory thresholds in AFD correlate with the cultivation temperature memory when tested immediately after training (Figure 1E). To examine whether this threshold persists long enough to store behavioral memory, animals trained at 15°C, 20°C or 25°C (T_c_) were subsequently held for 30 minutes at 20°C (T_H_) (Figure 1A). We observed that when the cultivation temperature memory (T_c_) differed from just-experienced holding temperatures (T_H_), AFD sensory threshold responses adapted to the new holding temperature (Figure 1I), although animals still performed according to the cultivation temperature memory (Figure 1F-H). While our findings are consistent with reports that behavioral memory persists for hours (Biron et al., 2006; Chi et al., 2007; Hedgecock and Russell, 1975; Mohri et al., 2005; Ramot et al., 2008b) and that AFD adaptation occurs in minutes (Ramot et al., 2008a; Yu et al., 2014), our findings are inconsistent with the hypothesis that the cultivation temperature memory resides in the AFD thermosensory thresholds. Instead, our findings suggest that the sensory threshold in AFD represents the holding temperature the animal has recently experienced.

How fast are AFD sensory thresholds adapting to newly experienced temperatures? We observed that AFD adapted to the holding temperature within minutes (Figure 1J & Figure S1B’-D’). AFD adaptation kinetics were mostly (>90%) complete within two minutes to increasing temperatures, and within thirty minutes to decreasing temperatures (Figure 1J). Analyses of AFD thermosensory response kinetics with distinct thermal stimuli (Figure S1H-S) and differing holding temperatures (Figure S1T-A’) further demonstrate that AFD responds to increases in temperature above the just-experienced ambient temperature (i.e., the holding temperature) without reference to the absolute temperature or the behaviorally preferred temperature (i.e., the cultivation temperature). Consistent with these observations, kinematic analyses of thermotaxis behavior indicate that animals continue to migrate towards their preferred cultivation temperature (T_c_) well after the AFD responses have adapted to T_H_ (Figure 1J,K), underscoring that the sensory threshold responses of AFD are separable from the cultivation temperature memory.

If the sensory thresholds in AFD are not the cultivation temperature memory, what do these sensory responses represent? Our findings indicate that the AFD thermosensory neuron detects temperature fluctuations from the recently-experienced ambient temperature (Figure 1 and Figure S1). The fast kinetics of adaptation allows AFD to maintain sensitivity to temperature changes while migrating in a gradient. This interpretation of the AFD thermosensory responses is consistent with observed kinetics of adaptation in calcium imaging (Yu et al., 2014) (Figure 1J and Figure S1B’-D’) and electrophysiological studies (Ramot et al., 2008a) performed in AFD.

We also note that this interpretation of AFD as an adaptable compass, and the kinetics of AFD adaptation, helps explain and unify previous observations in the field. For example, delays in thermotaxis performance based on cultivation temperature preference have been previously observed, and the nature of the differences discussed in the context of the cultivation temperature memory and the circuitry (Ito et al., 2006). Our findings predict these delays in the initiation of thermotaxis behavior based solely on the kinetics of AFD adaptation (Figure 1J,K). Moreover, the AFD adaptation kinetics also allowed us to predict and model AFD thresholds during behavior (Figure S1E’, dashed lines), highlighting that adaptation occurs prior to movement of the animals in the thermotaxis gradients (Figure S1E’, solid lines). Together, our findings indicate that sensory adaptation enables AFD to act as a compass, rendering the neuron maximally sensitive to local temperature variations and enabling it to encode thermosensory information as changes in temperature.

Next, we examined how AFD transfers thermal information onto its only chemical postsynaptic partner, AIY (White et al., 1986). We observed various temperature-dependent dynamics in AIY, but one distinct response was dependent on AFD: a response to warming near holding temperature (Clark et al., 2006) (Figure 2B,C and Figure S2). Consistent with this dependence, we observed that the AIY response to warming above holding temperature is similar in terms of timing to the responses seen in AFD. However, the frequency of animals displaying responses in AIY is lower than the frequency of animals displaying responses in AFD (Figure 2D-I and Figure S2).

**Figure 2.**
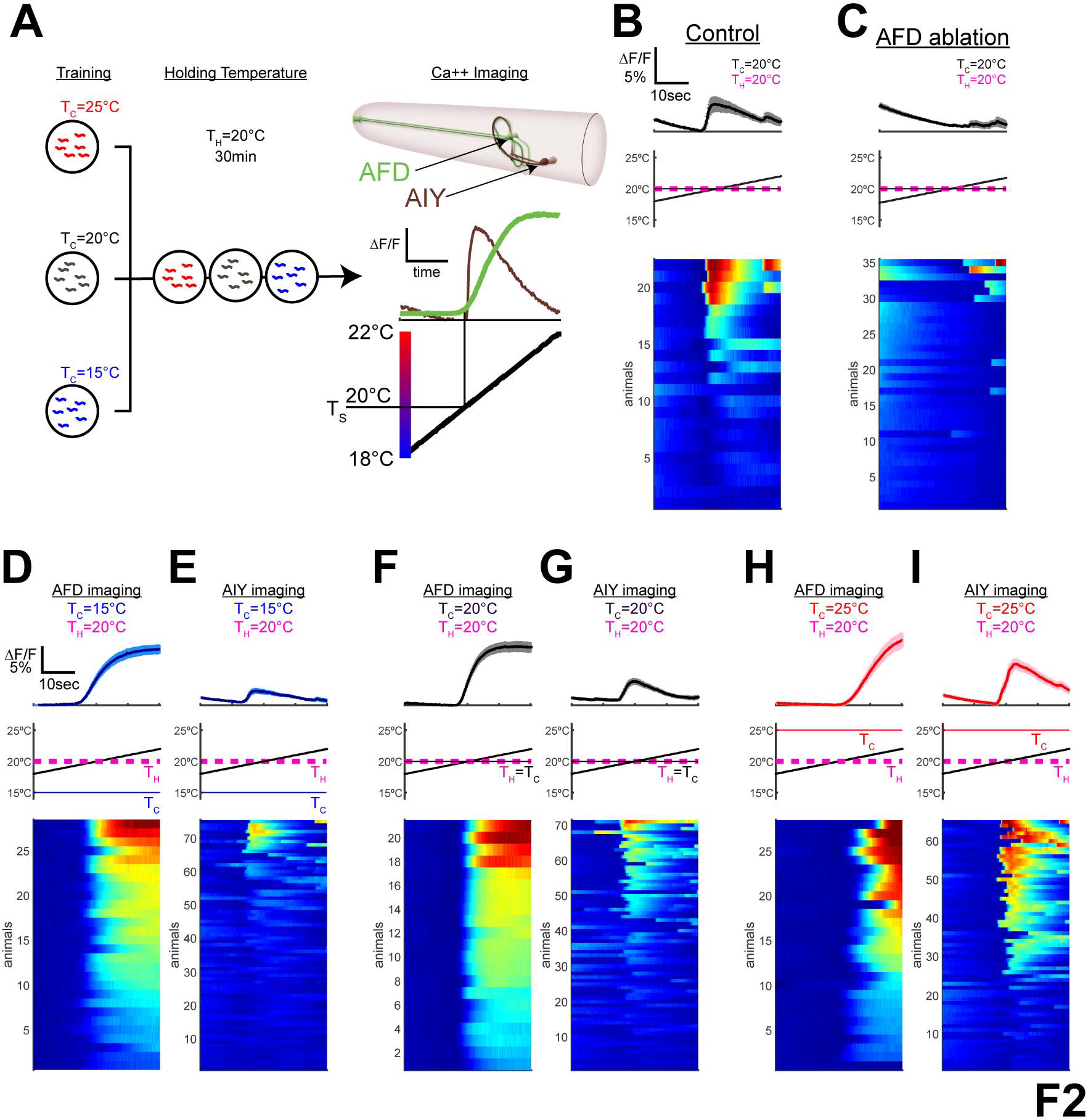
Postsynaptic response in AIY correlates with the cultivation temperature memory. (**A**) Schematic illustration of the experimental paradigm and calcium imaging in AFD (green) and AFD’s only chemical postsynaptic partner, AIY (brown). (**B,C**) Calcium dynamics of single AIY neurons in wild type in **B** or AFD-ablated animals in **C**; see also Figure S2. (**D-I**) Calcium dynamics of single AFD neurons (**D,F** and **H**) or AIY neurons (**E,G** and **I**) of animals trained at T_C_=15°C in **D** and **E**, T_C_=20°C in **F** and **G**, or T_C_=25°C in **H** and **I**, all with T_H_=20°C. Note that while AFD responds robustly in all conditions, the response frequency and amplitude in AIY varies depending on the cultivation temperature memory (see Figure S3 for quantification). Error bars denote SEM.

To better understand the relationship between the responses in AFD and the responses in AIY, we examined animals expressing GCAMP6 in AFD or AIY at varying cultivation temperatures (Figure 2). We observed that while warming above the holding temperature (20°C) reliably elicits responses in AFD regardless of T_C_ experience, the frequency of warming responses in AIY depends on the relationship between T_H_ and T_C_. In conditions in which T_H_>T_C_, in which wild type animals perform negative thermotaxis (i.e., move towards colder temperatures), AIY responses were infrequent (27/75 animals, ~36%). In conditions in which T_H_<T_C_, in which wild type animals perform positive thermotaxis (i.e., move towards warmer temperatures), AIY responses were frequent (44/65, ~68%). Intermediate response frequency (40/71, ~56%) was observed for AIY when T_H_=T_C_. Our findings are consistent with observations that AFD has an excitatory output onto AIY (Clark et al., 2006; Narayan et al., 2011) and suggest that cultivation temperature experience modulates this signal.

To understand how temperature preference is represented in the AIY responses, we examined mutants that affect temperature preference behaviors. Mutations that impair PKC-1 function do not affect AFD sensory responses (Luo et al., 2014) (Figure 3D,F,H and Figure S3A-F,M-O), yet produce robust thermotaxis behavioral defects (Okochi et al., 2005) (Figure 3A-C). PKC-1 loss-of-function mutants perform experience-independent positive thermotaxis, while expression of a constitutively active form of PKC-1 cell-specifically in AFD (*pAFD::caPKC-1*) produces experience-independent negative thermotaxis (Figure 3A-C). Defects observed with PKC-1 loss-of-function can be rescued by expression of wild-type PKC-1 in AFD alone (Okochi et al., 2005). Thus, both of positive and negative thermotaxis behaviors are dependent on PKC-1 activity specifically in AFD.

**Figure 3.**
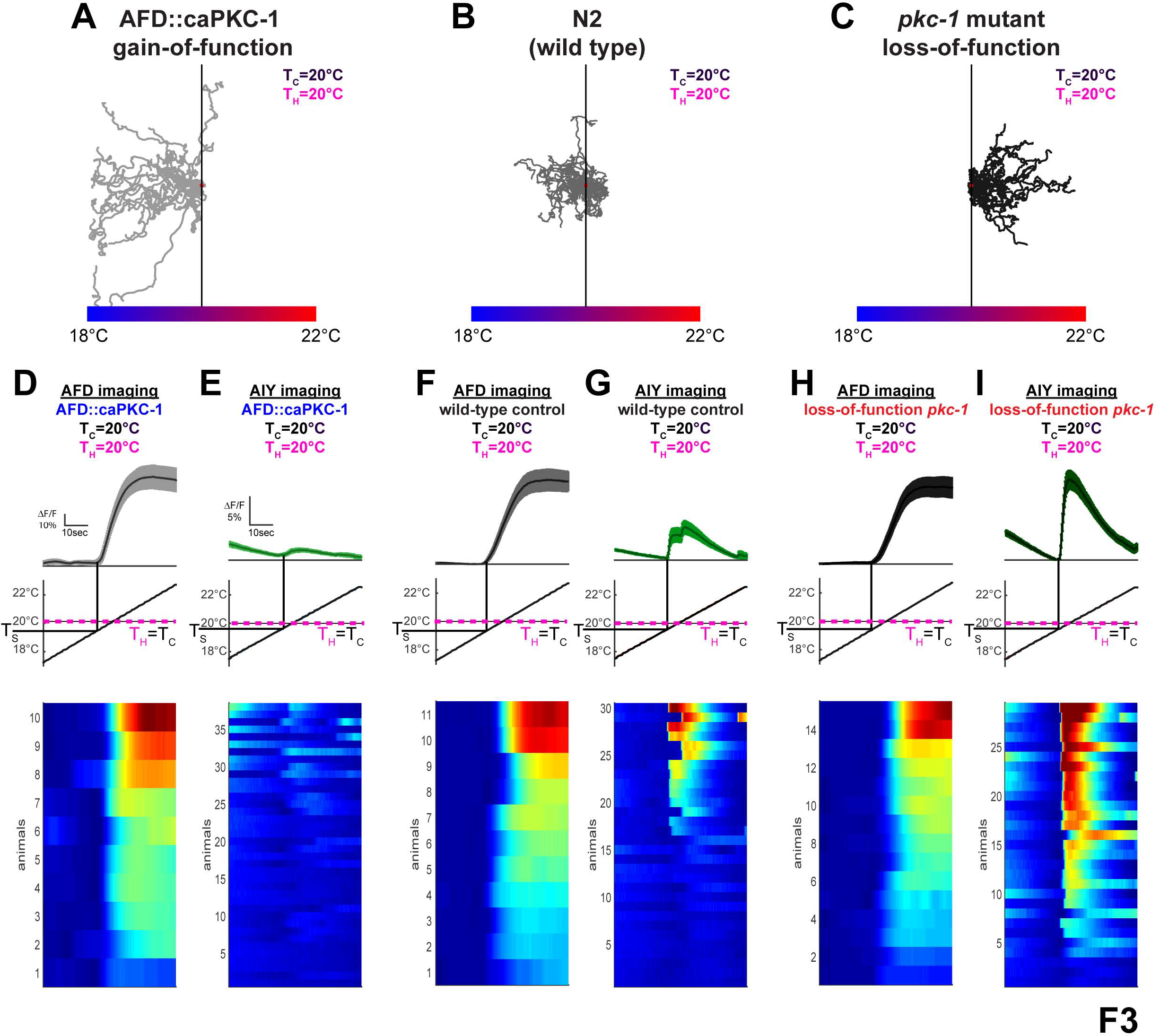
PKC-1 acts in AFD to control AIY response probability and T_C_ memory. (**A-C**) PKC-1 in AFD affects temperature preference regardless of previous experience. Expression of a constitutively active form of PKC-1 in the sensory neuron AFD produces worms that constitutively perform negative thermotaxis (**A**) while worms with a loss-of-function mutation in PKC-1 constitutively perform positive thermotaxis (**C**) regardless of T_c_ experience (Okochi et al., 2005). Wild-type worms T_C_=20°C show little bias when placed at the training temperature (**B**). (**D-I**) Calcium dynamics of single AFD neurons (**D, F**, and **H**) or AIY neurons (**E, G**, and **I**) for PKC-1 gain-of-function expressed specifically in AFD (AFD::caPKC-1) (**D** and **E**), wild type (**F** and **G**) or PKC-1 loss of function mutants (**H** and **I**) with T_C_= T_H_=20°C. Note how responses in AFD are not affected by PKC-1 genetic manipulations (consistent with earlier observations (Luo et al., 2014), see also Figure S3). In contrast, PKC-1 state in AFD alters AIY responses to phenocopy experiences that produce positive and negative thermotaxis. Error bars denote SEM.

When we examined temperature-dependent calcium responses in AIY in these genetic backgrounds, we observed that in worms expressing *pAFD::caPKC-1*, which perform experience-independent negative thermotaxis (Figure 3A), the frequency of the warming-evoked response in AIY (Figure 3E) is diminished as compared to wild type (Figure 3G) in similar conditions (Figure S3L). This genetic manipulation phenocopies the change in AIY calcium responses observed for wild type animals trained to perform negative thermotaxis (Figure 2E). Conversely, in the *pkc-1* loss-of-function mutants, which perform experience-independent positive thermotaxis (Figure 3C), warming response frequency in AIY increases (Figure 3I) relative to *pkc-1* loss-of-function (Figure 3E) and wild-type worms (Figure 3G) in similar conditions (Figure S3L), but phenocopies wild type animals trained to perform positive thermotaxis (Figure 3I). Our findings indicate that AIY thermal responses in *pkc-1* gain-of-function and loss-of-function mutants phenocopy those seen for wild-type animals trained to perform negative or positive thermotaxis, respectively. Our data also suggest that AIY response fidelity within a population is predictive of the temperature preference behavior. Importantly, our data suggest that PKC-1 regulates thermal preference by modulating presynaptic plasticity at the AFD to AIY synapse (Figure 3 and Figure S3).

Our findings support a model in which a synaptic gate from AFD to AIY, mediated by AFD presynaptic plasticity, determines the temperature preference and instructs whether worms respond to AFD-elicited temperature responses by moving up or down the gradient towards their previously learned preferred temperature. In animals that prefer warmer temperatures (or in animals lacking PKC-1), AFD responses to temperature increases are transduced to AIY, instructing positive thermotaxis. Conversely, in animals that prefer colder temperatures (or in animals with a PKC-1 gain of function in AFD) AFD responses to temperature increases are not transduced to AIY, and this “closed” synaptic gate is necessary to permit negative thermotaxis. Our results and model are also consistent with the long known effect of AIY inactivation on thermotaxis, which results in constitutively negative thermotaxis regardless of previous experience (Biron et al., 2006; Hobert et al., 1997; Kuhara et al., 2011; Luo et al., 2014; Mori and Ohshima, 1995).

If our model is correct, and expression of the cultivation temperature memory depends on regulation of synaptic communication between AFD and AIY, then directly coupling AIY to AFD via a synthetic electrical synapse would make animals perform positive thermotaxis regardless of previous experience. To test this prediction, we ectopically expressed the mammalian gap junction protein connexin 36 (CX36) (Rabinowitch et al., 2014) in both AFD and AIY. Expression of CX36 in either AFD or in AIY alone did not impact thermotaxis behavior in these animals (Figure S4A,B). However, expression of connexin channels in both AFD and AIY (*pAFD/pAIY::CX36*) enhanced AFD-elicited AIY responses (Figure 4B) without altering AFD thermosensory responses (Figure S4A-J). Specifically, we observed that animals trained to perform negative thermotaxis (T_H_>T_C_) displayed enhanced thermosensory responses in AIY when expressing the synthetic AFD:AIY gap junction (Figure 4B). Consistent with our model, these animals with the synthetic electrical synapses perform positive thermotaxis (Figure 4D) despite an experience history (T_H_>T_C_) that promotes negative thermotaxis (Figure 4C). These findings indicate that the synthetic electrical synapse bypasses endogenous plasticity of the AFD-AIY chemical synapse to recode the behavioral preference.

**Figure 4.**
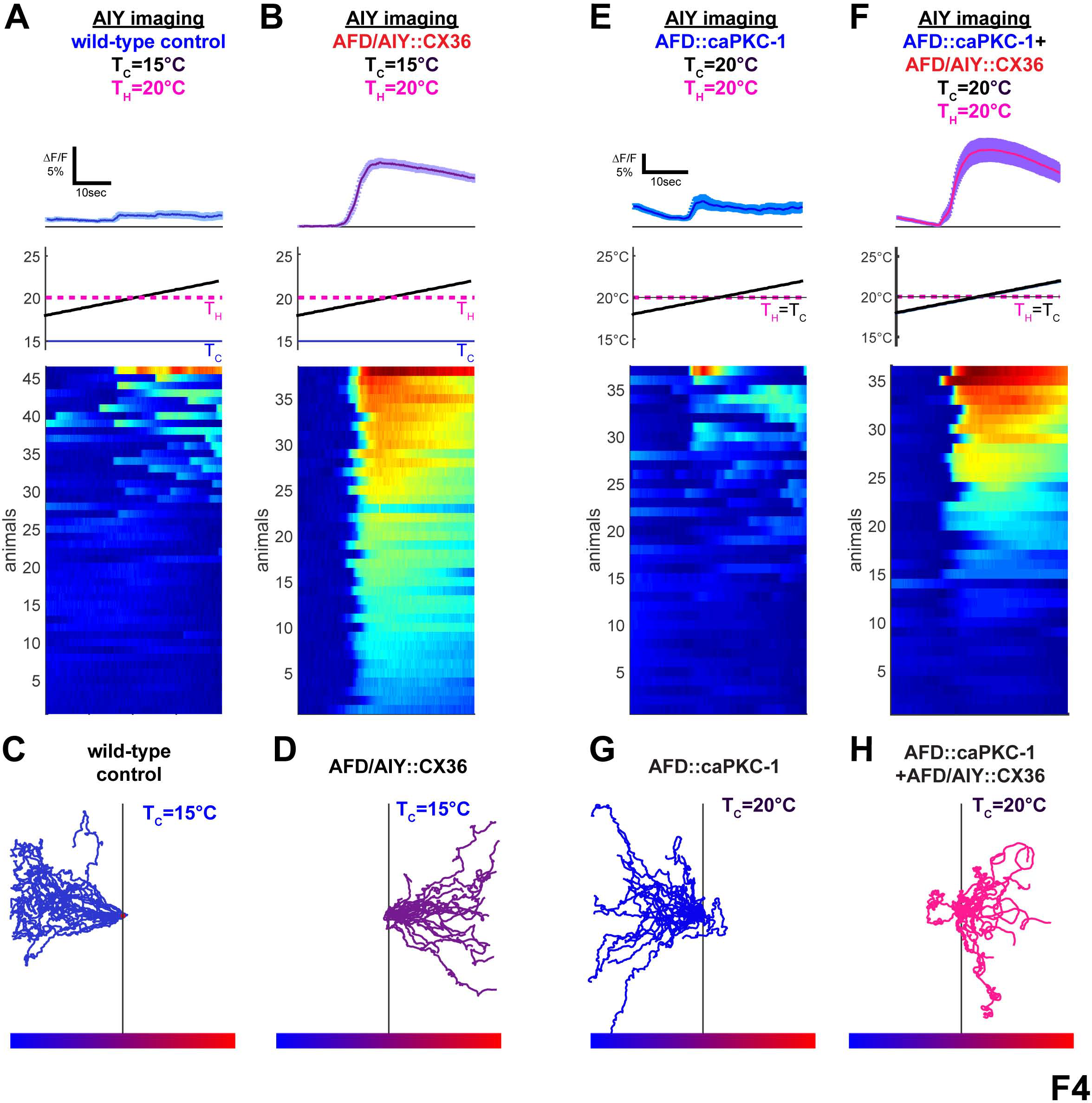
AFD to AIY connectivity actuates the thermal preference behavior. (**A**) Wild-type animals tested after adaptation above cultivation temperature (T_C_=15°C< T_H_=20°C) have weak and infrequent AIY responses near T_H_ (9/46 response penetrance, T_S_=19.0°C +/−0.7°C). (**B**) Under the same training conditions, ectopic expression of the gap junction protein CX36 in AFD and AIY (AFD/AIY::CX36) produces responses in AIY that mirror AFD responses (38/38 response penetrance, T_S_=19.0°C +/−0.2°C). (**C**) Worms cultivated at T_C_=15°C perform negative thermotaxis on a shallow thermal gradient centered at 20°C. (**D**) Expression of the gap junction protein CX36 in both AFD and AIY produces worms that perform positive thermotaxis under the same conditions. (**E**) Expression of a constitutively active form of PKC-1 in AFD dramatically diminishes AIY response frequency when T_C_=T_H_=20°C (13/37 response penetrance, T_S_=19.2°C +/−0.5°C). (**F**) AFD/AIY::CX36 increases AIY response frequency even in the presence of the AFD::caPKC-1 transgene (24/36 response penetrance, T_S_=19.3°C +/−0.2°C). Because of genetic linkage between the AFD::caPKC-1 transgene and the AFD::CX36 transgene, experiments in (**F** and **H**) were performed with unstable extrachrosomal lines as described in the Methods (unlike experiments in **B** and **D**, which were performed with an integrated CX36 transgenes). (**G-H**) In contrast to worms expressing constitutively active PKC-1 in AFD, which perform negative thermotaxis when T_C_=20°C (**G**), expression of CX36 in AFD and AIY (**H**) suppresses the phenotype of AFD::caPKC-1 animals and results in positive thermotaxis. For all experiments with CX36, see Figure S4 for corresponding controls of expression of hemichannels and AFD thermosensory responses. Error bars denote SEM.

We then examined whether the synthetic gap junction could bypass regulation of AFD-AIY plasticity by PKC-1. Consistent with our model, we observed that attenuation of AIY responses seen in the *pAFD::caPKC-1* animals (Figure 4E) is suppressed by expression of the synthetic AFD:AIY gap junction (Figure 4F). Moreover, the synthetic AFD:AIY gap junction was sufficient to convert negative thermotaxis observed in *pAFD::caPKC-1* mutants (Figure 4G) into positive thermotaxis (Figure 4H). Our findings indicate that modulation of the AFD:AIY synapse governs expression of the cultivation temperature memory.

## Discussion

The present work reconciles the observed fast timescale of AFD sensory adaptation with the hours-long behavioral memory and suggests that sensory adaptation and presynaptic plasticity, two independent plasticity modules, act cell-autonomously in AFD to actuate thermotaxis behavior. Our findings are also consistent with recent studies on AFD responses in freely moving animals navigating a gradient which demonstrate that AFD activities contain enough information to represent the thermal environment (Tsukada et al., 2016). Our findings extend those observations and demonstrate that AFD sensory responses are not dependent on the cultivation temperature memory, nor do they represent absolute temperatures (Clark et al., 2006; Kimura et al., 2004; Ramot et al., 2008a). Instead AFD responds to changes from the current temperature. We propose that AFD sensory responses act as an adaptable compass, making itself most sensitive to the local temperature environment to detect gradients. Together with electrophysiological studies (Ramot et al., 2008a), our findings suggest that warming and cooling might activate and inactivate AFD, respectively, allowing the temperature-evoked pattern of AFD activity to encode the direction of a worm’s movement in a temperature gradient.

The fast kinetics of AFD sensory adaption are likely to be important for maintaining perceptual constancy in a dynamic and sensory-rich environment. One of the better understood sensory adaptation systems is the vertebrate photoreceptor, which act as contrast detectors over wide dynamic ranges of illumination (Fain et al., 2001). Sensory adaptability in photoreceptor cells depends on a conserved mechanism that includes Guanylate Cyclases (GCs), Guanylate Cyclases Activating Protein (GCAP) and cGMP (Fesenko et al., 1985; Yau and Nakatani, 1985). Interestingly, molecular genetic studies in AFD have identified cGMP-dependent mechanism as necessary for temperature sensing (Inada et al., 2006; Komatsu et al., 1996; Ramot et al., 2008a; Wang et al., 2013), and recent studies in vertebrates and *C. elegans* suggest that guanylate cyclases respond to temperature (Chao et al., 2015; Takeishi et al., 2016). Our findings extend these studies, now demonstrating that sensory adaptation in AFD is important for thermotaxis behavior by enabling the animal to maintain contrast sensitivity and perceptual constancy over wide environmental variation.

Our study also demonstrates that presynaptic plasticity mechanisms contribute to the thermotaxis behavior by conferring the temperature preference. Modulation of temperature preference is actuated by PKC-1, which is required cell-autonomously in the presynaptic AFD neuron. PKC-1 is modulated by DAG, and consistent with our findings, a DAG regulator *dgk-3* has been shown to act cell-autonomously in AFD to control temperature preference without affecting AFD thermosensory responses (Biron et al., 2006). PKC-1 is the *C. elegans* homologue of nPKCε, a conserved molecule which has been implicated in presynaptic plasticity in the mammalian hippocampus and in memory (Son et al., 1996; Zisopoulou et al., 2013). Therefore, our studies suggest that a conserved nPKCε-dependent presynaptic plasticity mechanism acts in AFD to actuate the temperature preference memory through the control of an AFD to AIY synaptic gate. Our findings are consistent with the function of PKC-1 within other sensory neurons as a switch to reverse behavioral preference (Adachi et al., 2010; Tsunozaki et al., 2008) in a manner suggestive of presynaptic gating (Kunitomo et al., 2013; Tsunozaki et al., 2008). We extend these findings and demonstrate that the state of this AFD-AIY synaptic gate encodes whether AFD-derived thermosensory information will be interpreted as attractive or repulsive, depending on whether the worm is above or below its preferred temperature. On temperature gradients, AFD responds to increases in temperature regardless of the cultivation temperature memory. Positive thermotaxis occurs when these AFD responses are transmitted onto AIY, while negative thermotaxis occurs when these AFD responses fail to activate AIY.

Actuation of a learned preference behavior requires perceptual constancy during navigation of the sensory rich environments and the flexibility to differentially respond to sensory stimuli based on previous experiences. In this work, we have found that a two component system in a single sensory neuron is capable of encoding sensory-rich information, maintaining perceptual constancy, and flexibly assigning value based on memory. Bipartite plasticity through adaptation and synaptic mechanisms has been observed in at least one other *C. elegans* sensory neuron (Cho et al., 2016) and in Drosophila, where starvation produces complementary presynaptic facilitation and inhibition of sensory transmission to drive appetitive behavior (Ko et al., 2015; Root et al., 2011). Together, these findings suggest that the logical architecture composed of a directional sensory signal, integrated with a preference signal at the synapse, may be a common feature within sensory physiology. Therefore integration of plasticity modules, in the case of *C. elegans* thermotaxis within a single neuron, can cooperatively transform sensory information into a behavioral decision to actuate learned behavioral preference.

## Acknowledgements

We thank members of the Colón-Ramos lab and the Samuel lab for their thoughtful comments on the project. We also thank Miriam Goodman (Stanford University), Damon Clark (Yale University) and Piali Sengupta (Brandeis University) for thoughtful comments on the project and the manuscript. We thank the *Caenorhabditis* Genetics Center (supported by the National Institutes of Health (NIH), Office of Research Infrastructure Programs; P40 OD010440) for strains, and the Bargmann lab (Rockefeller University), the Goodman lab (Stanford University), the Schafer lab (Medical Research Council) and the Looger lab (Janelia Research Campus) for reagents. We thank Z. Altun (www.wormatlas.org) for diagrams used in figures. We thank the Research Center for Minority Institutions program and the Instituto de Neurobiología de la Universidad de Puerto Rico for providing a meeting and brainstorming platform. We thank the students of the Neural Systems and Behavior course from Marine Biological Laboratories at Woods Hole, particularly Ranran “Annette” Li and Raina Rhoades. This work was partially conducted at the Marine Biological Laboratories at Woods Hole, under a Whitman research award to D.A.C.-R. J.D.H was supported by the Ruth L. Kirschstein NRSA (NIH-NIMH-F32MH105063). Research in the Samuel laboratory was supported by National Institutes of Health (8DP1GM105383 and 1P01GM103770) and National Science Foundation (PHY-0957185. Research in the Colón-Ramos laboratory was also supported by NIH (R01NS076558) and the National Science Foundation (NSF IOS 1353845).

## Author Contributions

J.D.H and D.A.C.-R. designed experiments. J.D.H., A.C.C, A.A.P, M.L.T.S, I.R., and N.C. performed experiments and data analyses. J.D.H., A.C.C., A.A.P, M.L.T.S, N.C., L.L. and J.G. developed reagents and protocols. J.D.H., A.A.P., A.D.T.S, and D.A.C.-R. prepared the manuscript.

## Author Information

The authors declare no competing financial interests. Correspondence and request for materials should be addressed to D.A.C-.R.

## Materials and Methods

### Strains

Worms were cultivated at 20°C on nematode growth medium seeded with a lawn of *Escherichia coli* strain OP50 using standard methods (Brenner, 1974). The following strains were used in this study:

GN71 *pkc-1(pg5)*.
IK105 *pkc-1(nj1)*,
DCR3558 *olaIs23 [Pgcy-8::caPKC-1; Pgcy-8::tagRFP; Punc-122::dsRed]*,
DCR3055 *wyIs629 [Pgcy-8::GCaMP6s; Pgcy-8::mCherry; Punc-122::GFP]*,
DCR3245 *pkc-1(pg5); wyIs629 [Pgcy-8::GCaMP6s; Pgcy-8::mCherry; Punc-122::GFP]*,
DCR5099 *olaIs23 [Pgcy-8::caPKC-1; Pgcy-8::tagRFP; Punc-122::dsRed]; wyIs629 [Pgcy-8::GCaMP6s; Pgcy-8::mCherry; Punc-122::GFP]*,
DCR3056 *olaIs17 [Pmod-1::GCaMP6s; Pttx-3::mCherry; Punc-122::dsRed]*,
DCR3203 *pkc-1(pg5); olaIs17 [Pmod-1::GCaMP6s; Pttx-3::mCherry; Punc-122::dsRed]*,
DCR3572 *olaIs23 [Pgcy-8::caPKC-1; Pgcy-8::tagRFP; Punc-122::dsRed]; olaIs17 [Pmod-1::GCaMP6s; Pttx-3::mCherry; Punc-122::dsRed]*,
DCR3670 *olaIs17 [Pmod-1::GCaMP6s; Pttx-3::mCherry; Punc-122::dsRed]; olaIs23 [Pgcy-8::caPKC-1; Pgcy-8::tagRFP; Punc-122::dsRed]; olaEx2125 [Pgcy-8::caspase-3(p12)::nz; Pgcy-8::cz::caspase-3(p17); Pgcy-8::tagRFP; Pmyo-3::RFP]*,
DCR5586 *olaIs70 [Pgcy-8::CX36::mCherry; Pelt-7::GFP::NLS]*,
DCR5639 *olaIs72 [Pttx-3::sl2::CX36::mCherry; Pelt-7::mCherry::NLS]*,
DCR5793 *olaIs17 [Pmod-1::GCaMP6s; Pttx-3::mCherry; Punc-122::dsRed]; olaIs70 [Pgcy-8::CX36::mCherry; Pelt-7::GFP::NLS]; olaIs72 [Pttx-3::sl2::CX36::mCherry; Pelt-7::mCherry::NLS]*,
DCR5792 *wyIs629 [Pgcy-8::GCaMP6s; Pgcy-8::mCherry; Punc-122::GFP]; olaIs70 [Pgcy-8::CX36::mCherry; Pelt-7::GFP::NLS]; olaIs72 [Pttx-3::sl2::CX36::mCherry; Pelt-7::mCherry::NLS]*,
DCR5789 *olaIs17 [Pmod-1::GCaMP6s; Pttx-3::mCherry; Punc-122::dsRed]; olaIs70 [Pgcy-8::CX36::mCherry; Pelt-7::GFP::NLS]*,
DCR5790 *olaIs17 [Pmod-1::GCaMP6s; Pttx-3::mCherry; Punc-122::dsRed]; olaIs72 [Pttx-3::sl2::CX36::mCherry; Pelt-7::mCherry::NLS]*,
DCR5404 *olaIs17 [Pmod-1::GCaMP6s; Pttx-3::mCherry; Punc-122::dsRed]; olaIs23 [Pgcy-8::caPKC-1; Pgcy-8::tagRFP; Punc-122::dsRed]; olaEx3219 [Pgcy-8::CX36::mCherry; Pttx-3::sl2::CX36::mCherry; Pmyo-3::RFP]*,
DCR5380 *olaIs17 [Pmod-1::GCaMP6s; Pttx-3::mCherry; Punc-122::dsRed]; olaEx3219 [Pgcy-8::CX36::mCherry; Pttx-3::sl2::CX36::mCherry; Pmyo-3::RFP]*

### Behavioral Assay

Animals were reared at 20°C for all experiments with shifts to the training temperature 4 hour prior testing. High-throughput behavioral analyses were performed as described (Luo et al., 2014). Briefly, synchronized young adult populations were washed in M9 (Stiernagle, 2006), then transferred by pipette to the 20°C isotherm of the behavioral test plate (22-cm x 22-cm agar plates with 18°C to 22°C thermal gradient). Migration was monitored for >30 min at 2fps using a MightEx camera (BCE-B050-U). Trajectories were analyzed using an adaptation of the MagatAnalyzer software package as previously described (Gershow et al., 2012; Luo et al., 2014).

### Molecular Biology and transgenic lines

Gateway recombination (Invitrogen) was used to generate *C. elegans* expression plasmids for caPKC-1, GCaMP6, caspases and CX36. For PCR-based cloning and subcloning of components into the Gateway system, either Phusion or Q5 High-Fidelity DNA-polymerase (NEB) was used for amplification. All components were sequenced within the respective Gateway entry vector prior to combining components into expression plasmids via a four-component Gateway system (Merritt and Seydoux, 2010). For production of the caPKC-1 transgene, wild-type *pkc-1* cDNA was isolated from a mixed stage *C. elegans* cDNA library using primers amplifying transcript F57F5.5b and overhangs that allowed subsequent recombination into the vector pDONR221 (Invitrogen). The pseudosubstrate domain mutation (A160E) (Okochi et al., 2005) was introduced using QuikChange mutagenesis (Stratagene). GCaMP6s, cz::caspase-3(p17), caspase-3(p12)::nz, and CX36 were introduced into pDONR221 using a similar PCR-based strategy from plasmid sources (Chelur and Chalfie, 2007; Chen et al., 2013; Rabinowitch et al., 2014). Cell-specific promoters were introduced using the pENTR 5’-TOPO vector (Invitrogen) after amplification from genomic DNA or precursor plasmids. Transgenic lines were created by microinjection into the distal gonad syncytium (Mello and Fire, 1995) and selected based on expression of one or more co-injection markers: Punc-122::GFP, Punc-122::dsRed, Pelt-7::GFP::NLS, Pelt-7::GFP::NLS, or Pmyo-3::RFP. In those cases where extrachromosomal arrays were integrated, incorporation into the genome was induced with trimethylpsoralen (TMP) activated by UV using standard procedures followed by outcross. All integrated lines were outcrossed at least four times before testing.

### Calcium Imaging

For imaging, worms were mounted on a thin pad of 5% agarose in M9 buffer between two standard microscope coverslips. Worms were immobilized with 7.5-10mM levamisol (Sigma). All solutions were pre-equilibrated to the holding temperature prior to sample preparation. Prepared coverslip assemblies were placed at TH on the peltier surface of the thermoelectric control system as illustrated in Figure S1A. A thermal probe (SRTD-2, Omega) mounted onto the surface of the peltier was used for feedback control via a commercially available proportional-integral-derivative (PID) controller (FTC200, Accuthermo). Target temperatures were supplied to the PID controller by a custom computer interface written in LabView (National Instruments), which subsequently gated current flow from a 12V power supply to the peltier via an H-bridge amplifier (FTX700D, Accuthermo). Excess heat was removed from the peltier with a water cooling system. Precise temperature control was initially confirmed with an independent T-type thermal probe (IT-24P, Physitemp) attached to a hand-held thermometer (HH66U, Omega) and routinely compared to incubator temperatures with an infrared temperature probe (Actron). After mounting worms and placing them on the thermal control stage, fluorescence time-lapse imaging was begun immediately prior to implementing the temperature protocol in LabView, and temperature readings were recorded continuously while imaging. Fluorescence time-lapse imaging (250ms exposure) was performed using a Leica DM5500 microscope with a 10X/0.40 HC PL APO air objective. A Photometrics Dual-View 2 (DV2) optical splitter was used to simultaneously project distinct green and red fluorescence images onto the two halves of a Hamamatsu ORCA-Flash4.0 LT camera. Image acquisition, segmentation into regions of interest, and color alignment were performed in MetaMorph (Molecular Devices). Downstream data processing was performed using custom scripts written in MATLAB (Mathworks), including alignment of fluorescence intensity values to the temperature stimulus, ratiometric signal analyses, calcium response detection, and initial figure production. Temperature readings were assigned to image frames in MATLAB based on CPU timestamps on images and temperature readings. We employed best practices in acquisition and analyses of calcium imaging data in live animals, including the use of stably integrated transgenic lines with wild type thermotaxis behaviors, robust analyses of large number of animals and blind scoring of phenotypes (Broussard et al., 2014; Chen et al., 2013; Tian et al., 2012). Our calcium imaging strain and microscope optical design allow ratiometric analyses comparing GCaMP6 green fluorescence to signal from a calcium-independent red fluorophore, mCherry. Comparing our analysis using ratiometric normalization with analysis using the green channel alone, we observed indistinguishable patterns of activity. Because of this observation and because some subsequently used markers (e.g. *Pmyo-3::RFP* strains) produce additional red fluorescent signal in the head that can alter ratio baselines, we report fluorescence for the green channel alone. Response thresholds were determined with both computer-assisted response selection by a blind human observer and an automated script utilizing signal intensity and its derivative; these two methods showed high concordance. Heat map scaling, response graphing, and response detection thresholds were performed with the same consistent settings within each experiment. Heat maps were displayed using the Matlab ‘imagesc’ function, which scales data to the full color range available. The same scaling was applied to all comparable data within a figure. Unless otherwise specified, the scaling was based upon the highest intensity data set to minimize saturation. For analyses of AFD calcium signals, we selected four regions of interest: sensory ending, soma, axon, and a control region outside of AFD. We observed similar response profiles for all three regions within AFD, consistent with previous reports (Clark et al., 2006). Because the sensory ending is isolated from other sources of fluorescence (such as autofluorescence) and is the identified origin of thermosensory signals (Clark et al., 2006), the reported responses in our manuscript are at the sensory endings. For analyses of AIY, we divided the neuron into four previously defined and morphologically distinct regions based on EM reconstruction micrographs (White et al., 1986), as described (Colon-Ramos et al., 2007): 1) soma; 2) an asynaptic zone proximal to the soma (Zone 1); 3) a synapse-rich ~5uM long region in which the neurite turns dorsally from the anterior ventral nerve cord into the nerve ring (Zone 2); and 4) a distal segment within the nerve ring with sparse presynaptic distribution (Zone 3). In our analyses, we quantified signal intensity in the AIY soma and synaptic Zone 2 and Zone 3 regions. We also quantified a reference region outside of AIY. Consistent with previous reports (Clark et al., 2006), we observed clear and qualitatively similar responses to thermal stimuli in the synaptic Zone 2 and Zone 3 regions. Although calcium responses were sometimes observed in the AIY soma, we could not correlate these soma responses to the thermal stimuli.

### Modeling of AFD adaptation

After training, AFD response threshold adapts to a new temperature with time following an exponential function (T_S_=C*e^−t/tau^, Figure 1J). The change in T_S_, ∆T_S_, with an increment of time, ∆t, at time t can be expressed as ∆T_S_/∆t(t)=(−C(t)/tau)*e^−t/tau^. The value of C at time t, C(t), is the offset between current temperature, T^H^(t), and AFD threshold, TS(t), such that C(t)=T_S_(t)-T_H_(t)). Based on these three equations and the empirically calculated time constants for tau (derived from data in Figure 1J), the AFD threshold can be calculated iteratively for a migrating animal by incrementing the T_S_ value with the derivative of T_S_ under present conditions, T_S_(t)=T_S_(t-1)+∆TS/∆t (t)* ∆t, resulting in the model presented in (Figure S1E’). Because tau is distinct for adaptation to higher and lower temperature, the appropriate tau was selected at each iteration depending on the relationship between T_S_ and T_H_.

### Statistical analyses

Continuous data were analyzed for statistical significance by non-parametric Kruskall-Wallis ANOVA followed by *post-hoc* comparisons with a two-tailed Wilcoxon rank sum test. Categorical data were analyzed for statistical significance by chi-squared analysis followed by pair-wise two-tailed Fisher’s exact tests.

**Figure S1.**
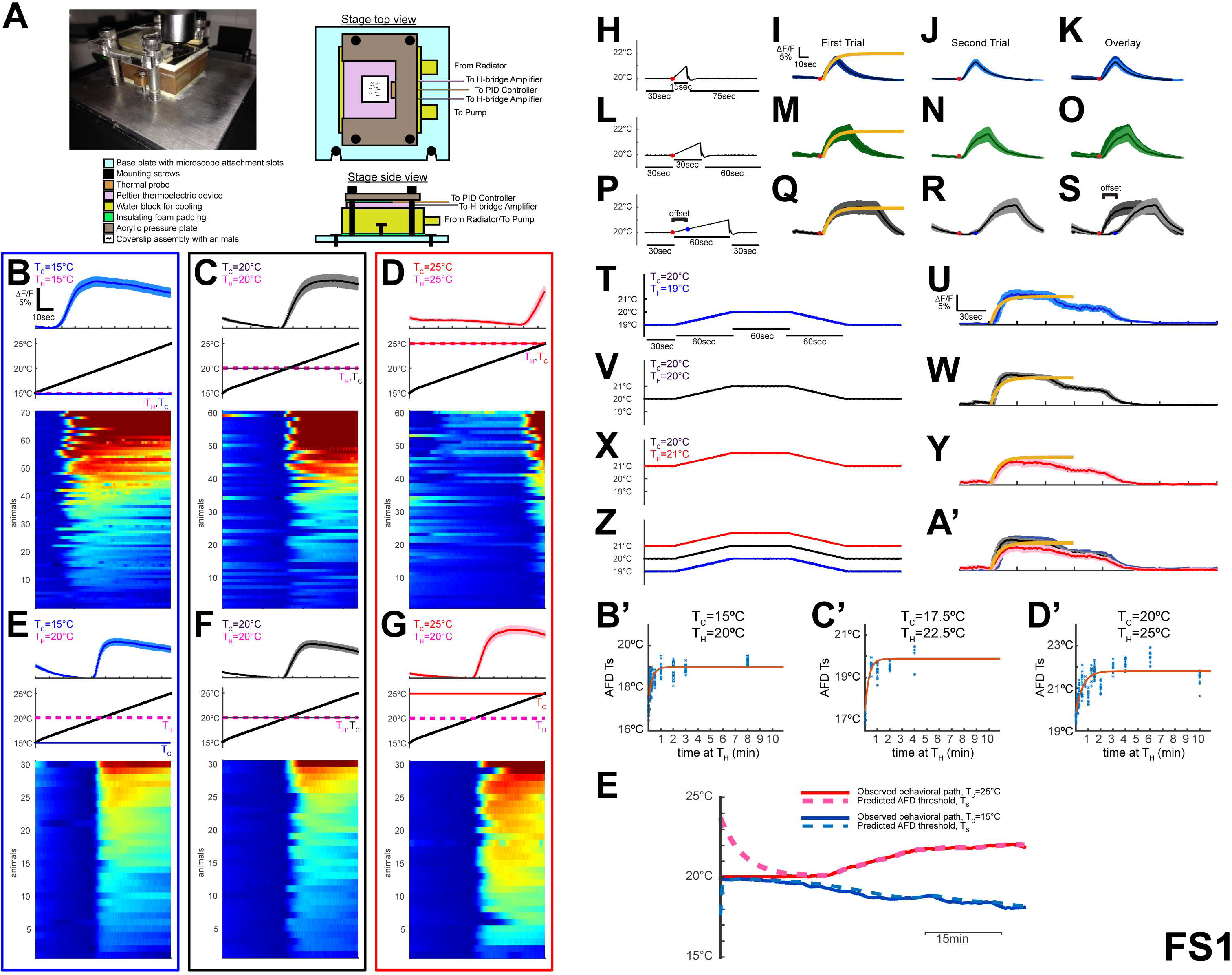
AFD thermosensory responses adapt rapidly and encode variations in temperature. Related to Figure 1. (**A**) Picture and schematic of the thermoelectric control system used for calcium imaging. Animals immobilized by levamisole were mounted on an agarose pad between two coverslips and placed directly onto the thermoelectric device. Precise temperature control at the peltier surface is achieved via a proportional-integral-derivative (PID) controller (Accuthermo) that gates applied current via an H-bridge amplifier (Accuthermo). A water-cooling system extracts excess heat, as represented in the schematic. Specific temperature protocols were supplied to the PID controller through a custom computer interface written in LabView (see Methods for additional information). (**B-D**) Calcium imaging of AFD responses to a rising linear ramp of 0.1°C/sec from 15°C to 25°C on animals trained at T_C_=15°C in **B**, T_C_=20°C in **C**, or T_C_=25°C in **D**. Mean responses (top panel), stimulus ramp (middle panel), and heat maps (bottom panel) are shown for each condition. Mean responses (dark lines) are illustrated with standard error represented by lighter shaded areas. Heat maps (bottom panel) illustrate the individual responses from >60 animals (each animal is a row) through time. Because of the large sample size in these panels and the variance in AFD response amplitude, some samples are illustrated as saturated signals to allow the frequency of response to be apparent in the heat map. Note that AFD thermosensory responses occur at a threshold temperature (T_S_) that adapts with experience for animals raised at T_C_=15°C (T_S_=16.9°C+/−0.5°C) in **B**, for animals raised at T_C_=20°C (T_S_=19.8°C+/−0.5°C) in **C**, and for animals raised at T_C_=25°C (T_S_=23.9°C+/−0.4°C) in **D**. The data shown here correspond to that of Figure 1E. (**E-G**) Like **B-D**, but after 30min of adaptation at 20°C (T_H_=20°C) as described in Figure 1A. Note that after 30min of T_H_=20°C, animals previously trained at T_C_=15°C, 20°C or 25°C show a response threshold very near T_H_ and not T_C_ (T_S_=19.4°C+/−0.1°C; T_S_=19.8°C+/−0.2°C and T_S_=19.9°C+/−0.3°C respectively). The data shown here correspond to that of Figure 1I. (**H-S**) AFD calcium responses indicate direction and duration of temperature change. Animals held at the cultivation temperature (T_C_=20°C) were then exposed to a defined thermal ramp of 1°C over 15sec in **H-K** (n=10), 30sec in **L-O** (n=8), or 60sec in **P-S** (n=11). Stimulus onset is marked with a red dot. For each stimulus protocol in **H, L**, and **P**, two immediately consecutive tests were performed on the same animals (first response in **I, M** and **Q** and second response in **J, N** and **R**). An overlay of the two consecutive responses are shown in **K, O** and **S**. The AFD calcium rise fits a single exponential function related to the duration of ramping stimulus (y=C*(1-e^−kt^), yellow curve in **I, M** and **Q**). Note that while overlay of first and second responses fit well in **K** and **O**, we observed a shift in the response timing (“offset”) of the second response with a 60sec stimulus in **R**, as illustrated in the overlay, **s**. The observed shift for this 60sec ramp is consistent with the rapid adaptation kinetics of the AFD threshold (Figure 1J), and suggest that the shift is due to adaptation experienced by the animal during the 60sec test stimulus of the first trial in **Q**. (**T-A’**) AFD responds to changes in temperature independently of absolute temperature, consistent with previous observations (Clark et al., 2006; Kimura et al., 2004) and independent of cultivation temperature memory. Animals cultivated at T_C_=20°C were held for 30min at T_H_=19°C in **U** (n=29), 20°C in **W** (n=26), or 21°C in **Y** (n=24), and then subjected to the protocols in **T, V** and **X**. An overlay of these responses in **A’** illustrates that AFD responds similarly regardless of holding temperatures, and that the rising phase of the response fits an exponential function (yellow curve in **I**, identical to curves in **M, Q, U, W, Y**, and **A’**). (**B’-D’**) AFD thermosensory thresholds adapt within minutes to new holding temperatures. Animals trained at T_C_=15°C in **B’** (n=269 total animals examined, with each dot representing a measurement from a single animal. 17-44 animals were examined for each timepoint, for a total of 9 timepoints as indicated in the graph), T_C_=17.5°C in **C’** (n=75 with 7-15 animals examined for each timepoint, for a total of 6 timepoints as indicated in the graph), or T_C_=20°C in **D’** (n=233 with 8-37 animals examined for each timepoint, for a total of 14 timepoints as indicated in the graph) were then shifted to a holding temperature (T_H_=T_C_+5°C) for defined durations of time (as indicated in the graph) to assess AFD sensory adaptation dynamics. The threshold temperature of AFD response (T_S_) is plotted as a function of the time held at the new temperature (T_H_=20°C). Note that similarly rapid adaptation kinetics were observed regardless of the initial T_C_. AFD adaptation with these temperature shifts can be adequately modeled with a single exponential function. Adaptation over larger temperature shifts have been previously shown to fit a double exponential function, suggesting different adaptation kinetics and mechanisms depending on magnitude of temperature shift and time (Yu et al., 2014). (**E’**) Based on the kinetics we uncovered for AFD adaptation (Figure 1J & Figure S1B’-D’), we modeled AFD adaptation during behavior. The observed migratory route of a single worm with T_C_=15°C (blue solid line) or T_C_=25°C (red solid line) is illustrated over a 60min thermotaxis behavioral run. Using an exponential function, we calculated the predicted AFD threshold value, T_S_, at each time point based on the current temperature and previous T_S_ value as illustrated by the dashed light blue (T_C_=15°C) and light red (T_C_=25°C) lines (further explained in Methods). Note that AFD adapts such that during migration, the threshold closely mirrors current temperature on a migratory path.

**Figure S2.**
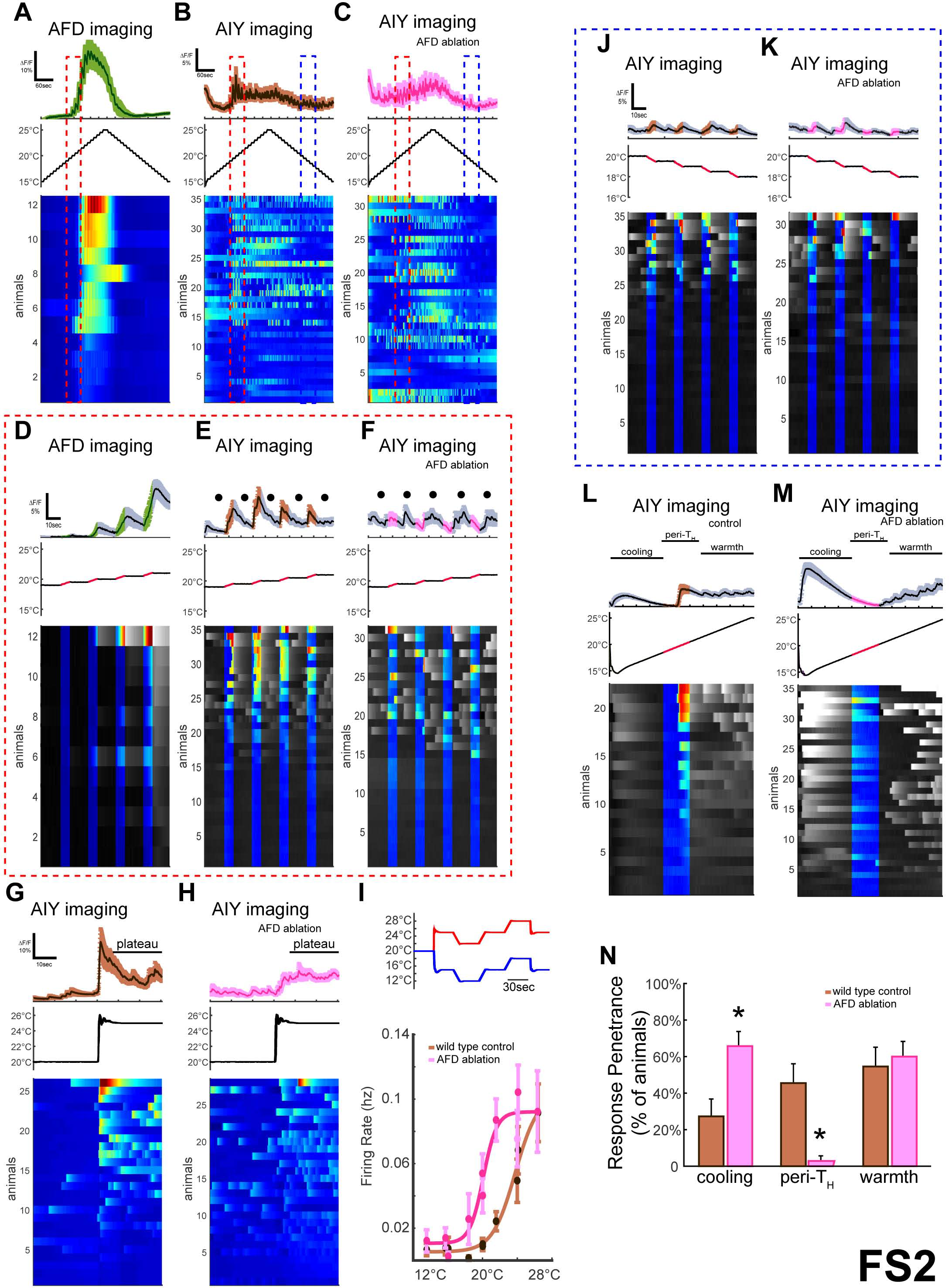
AIYs display distinct AFD-dependent and AFD-independent temperature responses. Related to Figure 2. (**A-C**) AFD in **A** and postsynaptic AIY in **B,C** calcium responses to thermal staircase protocol in wild type in **A,B** or AFD-ablated in **C** animals. In all panels, animals expressing GCaMP6 cell-specifically in AFD in **A** or AIY in **B,C** were trained at T_C_=20°C, anesthetized as described in Methods and examined at T_H_=20°C. Note that AFD responds only above a threshold near T_H_ (red dashed box) during the rising phase of the staircase protocol in **A**. AIY displays a complex pattern of responses to the staircase protocol, including synchronized responses on the ramping phase near the holding temperature (red dashed box in **B**). Cell-specific ablation of AFD alters the responses in AIY, most notably by eliminating the peri-T_H_ ramping response (red dashed box in **C**). (**D-F**) To highlight the responses eliminated by AFD ablation, the data from **A-C** are illustrated at higher magnification and ordered based on the magnitude of response during the peri-T_H_ ramping phase (red dashed box in **A-C**). In the top row, which represent the mean fluorescent traces for responses in AFD in **D** or AIY in **E,F** for the indicated stimuli (middle row), the plateau periods between temperature steps are in grayscale (and highlighted with black dots), whereas ramping periods are highlighted in color. Similarly, in the stimuli protocol schematized in the middle panels, ramps are highlighted in red and plateaus in black. Note that the onset of AFD response occurs during this ramping phase and at the temperature ramps in **A,D**, and that the AIY synchronized responses are also predominantly occurring at the temperature ramps in **E** (highlighted in brown in top panel; also highlighted in color in the calcium heat maps for individual animals in lower panels). Ablation of AFD does not eliminate all of AIY responses, but does abrogate the observed synchronized responses during the temperature ramps (Brown in **E** and red in **F** upper panels). Note also the emergence of ectopic response peaks during the plateau phases (black dots in mean trace of the upper panels). (**G-I**) Examination of AFD-independent plateau responses to temperature in AIY. Animals T_C_=20°C;T_H_=20°C were exposed to a single temperature step to 25°C and calcium responses were recorded in AIY for wild type in **G** and AFD-ablated in **H** animals. Note that the temperature increase elicits a synchronized response to temperature during the ramping phase in **G** and that this response is AFD-dependent in **H**. Also note that the synchronized response is followed by a period of calcium responses to the plateau period in which animals are held at the new temperature. Consistent with observations using protocols in **F**, these responses in AIY during the temperature plateau are not eliminated upon ablation of AFD in **H**. (**I**) Animals trained at T_C_=20°C; T_H_=20°C were subjected to one of the two protocols (red or blue) schematized in **I** (top panel). In these two protocols, animals are exposed to 30-sec holds either above (red) or below (blue) T_H_ as illustrated. Responses of mean firing frequency as a function of plateau temperature reveals a sigmoidal response curve with increased firing frequency in wild-type N2 worms above 20°C (brown curve, 26 animals were examined for each of the two protocols, for a total of 52 animals). In animals with AFD ablated (pink curve, n=28 animals were examined for the protocol above T_H_ and n=29 animals were examined for the protocol below T_H_), AIY responses during plateau periods are still observed with response frequency rising at lower temperatures than observed in control animals. These data suggest that the AIY plateau responses are not dependent on AFD, but are modulated by AFD. j, AIY also responds to temperature decrements as shown below 20°C of the staircase protocol from **A-C** (blue dashed box). (**K**) In animals with AFD ablated, AIY response peaks are still observed during temperature decrements below 20°C. (**L**) In worms with T_C_=20°C and T_H_=20°C, a single temperature decrement to 15°C followed by a linear ramp of 0.1°C/sec to 25°C illustrates responses to all three elements of the temperature protocol: decrement below 20°C (“cooling”, grayscale), warming near 20°C (“peri-TH”, color), and temperatures above 20°C (“warmth”, grayscale). (**M**) This protocol further illustrates that only the peri-T_H_ warming response is eliminated with AFD ablation. (**N**) Upon AFD ablation (pink), the percentage of animals observed to have an AIY calcium response (“penetrance”) selectively decreases during the peri-T_H_ window. Thus, AIY response frequency in the peri-T_H_ window is AFD dependent. Interestingly, a higher response penetrance can be noted in the cooling response after AFD ablation, consistent with AFD playing modulatory roles in these other temperature responses, similar to its modulatory role in the plateau responses to warming temperatures in **I**. * indicates p<0.05.

**Figure S3.**
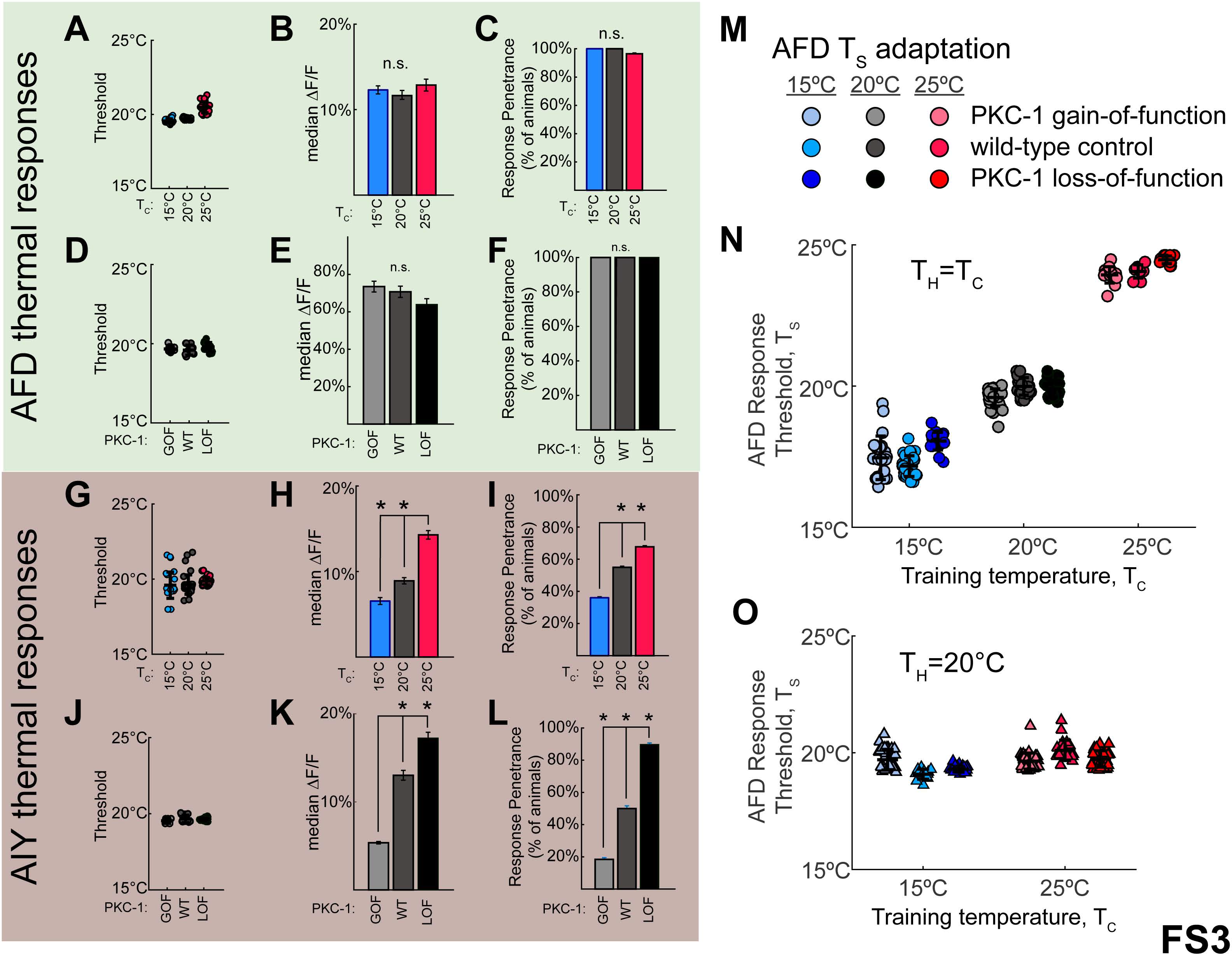
Directed migration on a thermal gradient is associated with changes in post-synaptic responses to AFD rather than changes in AFD sensory function. Related to Figures 2 and 3. (**A-C**) Animals cultivated at T_C_=15°C (blue, n=28), T_C_=20°C (gray, n=21), or T_C_=25°C (red, n=28) were exposed to T_H_=20°C for 30 minutes and subjected to a linear thermal ramp of 0.1°C/sec (as described in Figure 1A). Calcium imaging in AFD revealed that the response threshold values (T_S_) were very near T_H_=20°C for all conditions tested regardless of T_C_ (19.4°C +/−0.08°C, 19.6°C +/−0.05°C, and 20.2°C +/−0.26°C, respectively). We note that while animals migrating in a temperature gradient will experience temperature changes that can be up to 4°C different between the start point and the end point of the migration track (Figure 1K), the T_S_ values in this experiment approximate T_H_=20°C for all conditions, and that the value range varies less than 1.7°C between conditions. Moreover, no differences were observed in AFD responses for different T_C_ conditions, including median response amplitude in **B** or percentage of animals responding to the temperature ramps in **C**. (**D-F**) PKC-1 loss-of-function and gain-of-function mutants in AFD, which produce distinct thermal preference behaviors independent of experience (Okochi et al., 2005) (Figure 3A-C), were cultivated at T_C_=20° and examined for AFD calcium responses in linear thermal ramp protocols described in Figure 3D. Response threshold in **D**, median response amplitude in **E**, and response penetrance in **F** were quantified for animals with AFD-specific expression of PKC-1 gainof-function (“GOF”, light gray, n=10), wild-type animals (“WT”, gray, n=11), and genetic lesions resulting in PKC-1 loss-of function (“LOF”, black, n=15). Note that the PKC-1 genetic perturbations do not affect any of the observed AFD thermosensory responses, consistent with previous reports (Luo et al., 2014). (**G-I**) As **A-C**, but imaging calcium responses in AIY at T_C_=15°C (blue, n=75), T_C_=20°C (gray, n=71), or T_C_=25°C (red, n=65). Threshold for AIY response (the first observed response in the peri-T_H_ window) is similar regardless of T_C_ experience in **G**. Yet, the median response amplitude in **H** and the response penetrance in **I** in AIY vary depending on T_C_ experience. (**J-L**) As **D-F** but imaging calcium responses in AIY. No differences were observed in the AIY response threshold between the wild type and PKC-1 loss of function or gain of function mutants in **J**, but cold-seeking PKC-1 GOF animals (light gray, n=38) show significantly reduced mean response amplitude in **K** and response penetrance in **L** in comparison to wildtype (dark gray, n=30) and warmth-seeking PKC-1 LOF mutant animals (black, n=29). (**M-N**) AFD response thresholds for PKC-1 gain-of-function, wild type or PKC-1 loss-of-function mutants trained at different T_C_ as indicated in the legend in **M**. Each circle represents the response of a single animal. (**O**) From T_C_=15°C (blue) or T_C_=25°C (red), animals were then held at T_H_=20°C for 30min. For all three genotypes, T_S_ values adapted to nearly 20°C within this period. Together, these findings indicate that AFD response thresholds and adaptability are not impacted by in PKC-1 GOF or LOF mutants. * indicates p<0.05

**Figure S4.**
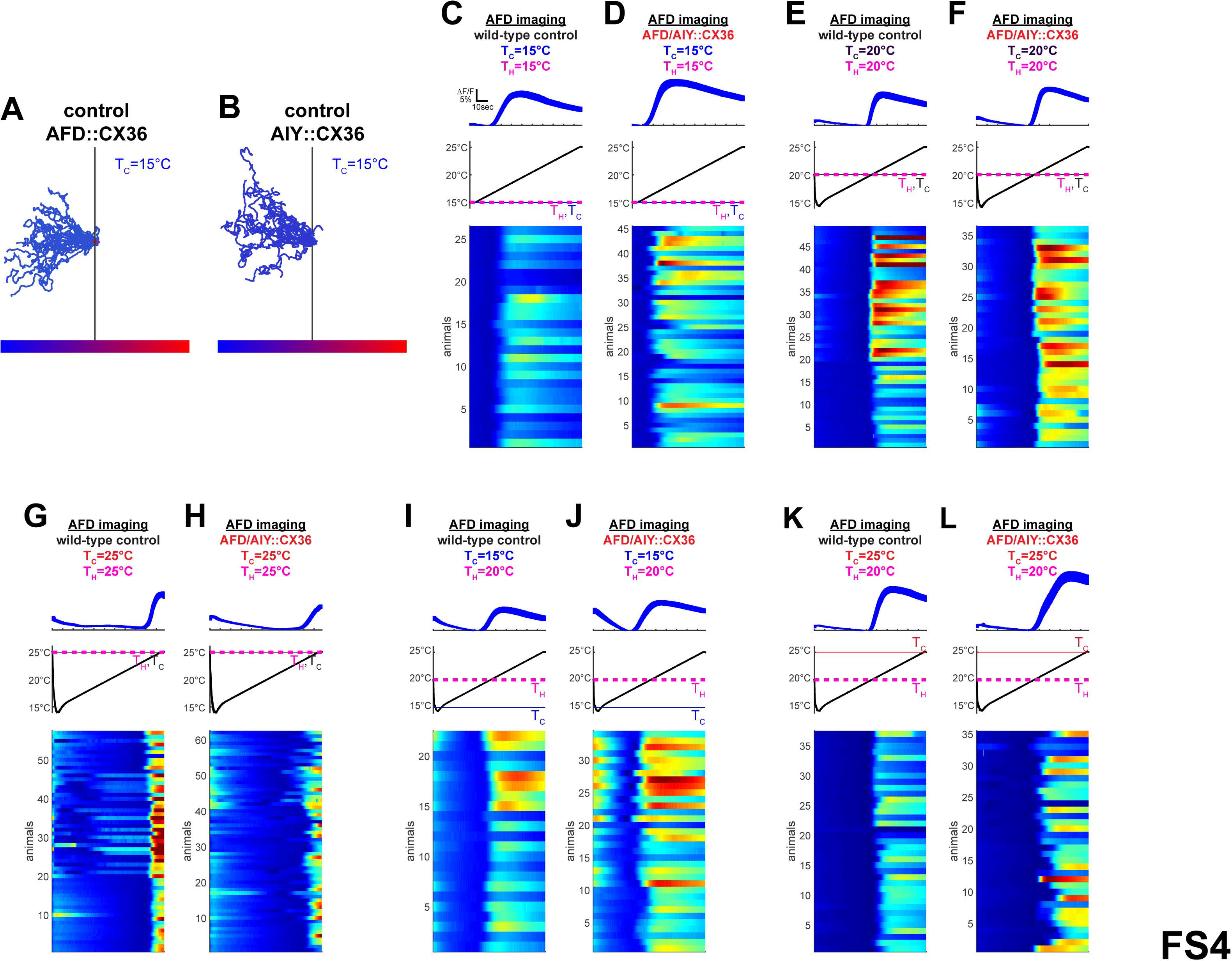
CX36 behavioral phenotype is not a consequence of hemi-channel formation or altered AFD sensory function. Related to Figure 4. (**A-B**) Stable, integrated lines expressing CX36 in individual AFD (**A**) or AIY (**B**) neurons were created and then examined in thermotaxis behavior. Note that expression of the CX36 hemichannel in either AFD or AIY does not affect thermotaxis behavior for animals with T_C_=15°C (n=25 for both experiments). These same integrated lines were then crossed to each other to generate the line examined in Figure 4B,D. (**C-H**) Expression of CX36 in AFD and AIY does not alter AFD sensory responses. In animals expressing CX36, AFD calcium signals were examined in response to a linear temperature ramp of 0.1°C/sec. When holding and cultivation temperatures were the same (T_H_=T_C_), wild-type animals (**C, E** and **G**) or animals containing the synthetic AFD to AIY gap junction (**D, F** and **H**) display similar responses in AFD. These findings suggest that the synthetic CX36 AFD:AIY gap junction does not affect AFD responses to temperature as compared to wild type. (**I-L**) AFD threshold adaptation was also examined for wild type (**I,K**) and mutant (**J,L**) animals containing the synthetic AFD to AIY gap junction with T_C_=15°C;T_H_=20°C (**I,J**) or T_C_=25°C;T_H_=20°C (**K,L**). Note that AFD adaptability is similar to wild type even upon expression of the synthetic gap junction.

## References

[1] Adachi, T., Kunitomo, H., Tomioka, M., Ohno, H., Okochi, Y., Mori, I., and Iino, Y. (2010). Reversal of salt preference is directed by the insulin/PI3K and Gq/PKC signaling in Caenorhabditis elegans. Genetics 186, 1309–1319.

[2] Aoki, I., and Mori, I. (2015). Molecular biology of thermosensory transduction in C. elegans. Curr Opin Neurobiol 34, 117–124.

[3] Basu, J., and Siegelbaum, S.A. (2015). The Corticohippocampal Circuit, Synaptic Plasticity, and Memory. Cold Spring Harb Perspect Biol 7.

[4] Beverly, M., Anbil, S., and Sengupta, P. (2011). Degeneracy and neuromodulation among thermosensory neurons contribute to robust thermosensory behaviors in Caenorhabditis elegans. The Journal of neuroscience: the official journal of the Society for Neuroscience 31, 11718–11727.

[5] Biron, D., Shibuya, M., Gabel, C., Wasserman, S.M., Clark, D.A., Brown, A., Sengupta, P., and Samuel, A.D. (2006). A diacylglycerol kinase modulates long-term thermotactic behavioral plasticity in C. elegans. Nature neuroscience 9, 1499–1505.

[6] Biron, D., Wasserman, S., Thomas, J.H., Samuel, A.D., and Sengupta, P. (2008). An olfactory neuron responds stochastically to temperature and modulates Caenorhabditis elegans thermotactic behavior. Proceedings of the National Academy of Sciences of the United States of America 105, 11002–11007.

[7] Chao, Y.C., Chen, C.C., Lin, Y.C., Breer, H., Fleischer, J., and Yang, R.B. (2015). Receptor guanylyl cyclase-G is a novel thermosensory protein activated by cool temperatures. EMBO J 34, 294–306.

[8] Chen, T.W., Wardill, T.J., Sun, Y., Pulver, S.R., Renninger, S.L., Baohan, A., Schreiter, E.R., Kerr, R.A., Orger, M.B., Jayaraman, V., et al. (2013). Ultrasensitive fluorescent proteins for imaging neuronal activity. Nature 499, 295–300.

[9] Chi, C.A., Clark, D.A., Lee, S., Biron, D., Luo, L., Gabel, C.V., Brown, J., Sengupta, P., and Samuel, A.D. (2007). Temperature and food mediate long-term thermotactic behavioral plasticity by association-independent mechanisms in C. elegans. J Exp Biol 210, 4043–4052.

[10] Cho, C.E., Brueggemann, C., L’Etoile, N.D., and Bargmann, C.I. (2016). Parallel encoding of sensory history and behavioral preference during Caenorhabditis elegans olfactory learning. Elife 5.

[11] Clark, D.A., Biron, D., Sengupta, P., and Samuel, A.D. (2006). The AFD sensory neurons encode multiple functions underlying thermotactic behavior in Caenorhabditis elegans. The Journal of neuroscience: the official journal of the Society for Neuroscience 26, 7444–7451.

[12] Davis, R.L. (2011). Traces of Drosophila memory. Neuron 70, 8–19.

[13] Fain, G.L., Matthews, H.R., Cornwall, M.C., and Koutalos, Y. (2001). Adaptation in vertebrate photoreceptors. Physiol Rev 81, 117–151.

[14] Fesenko, E.E., Kolesnikov, S.S., and Lyubarsky, A.L. (1985). Induction by cyclic GMP of cationic conductance in plasma membrane of retinal rod outer segment. Nature 313, 310–313.

[15] Garrity, P.A., Goodman, M.B., Samuel, A.D., and Sengupta, P. (2010). Running hot and cold: behavioral strategies, neural circuits, and the molecular machinery for thermotaxis in C. elegans and Drosophila. Genes Dev 24, 2365–2382.

[16] Gjorgjieva, J., Drion, G., and Marder, E. (2016). Computational implications of biophysical diversity and multiple timescales in neurons and synapses for circuit performance. Curr Opin Neurobiol 37, 44–52.

[17] Hedgecock, E.M., and Russell, R.L. (1975). Normal and mutant thermotaxis in the nematode Caenorhabditis elegans. Proceedings of the National Academy of Sciences of the United States of America 72, 4061–4065.

[18] Hobert, O., Mori, I., Yamashita, Y., Honda, H., Ohshima, Y., Liu, Y., and Ruvkun, G. (1997). Regulation of interneuron function in the C. elegans thermoregulatory pathway by the ttx-3 LIM homeobox gene. Neuron 19, 345–357.

[19] Inada, H., Ito, H., Satterlee, J., Sengupta, P., Matsumoto, K., and Mori, I. (2006). Identification of guanylyl cyclases that function in thermosensory neurons of Caenorhabditis elegans. Genetics 172, 2239–2252.

[20] Ito, H., Inada, H., and Mori, I. (2006). Quantitative analysis of thermotaxis in the nematode Caenorhabditis elegans. J Neurosci Methods 154, 45–52.

[21] Kimura, K.D., Miyawaki, A., Matsumoto, K., and Mori, I. (2004). The C. elegans thermosensory neuron AFD responds to warming. Curr Biol 14, 1291–1295.

[22] Ko, K.I., Root, C.M., Lindsay, S.A., Zaninovich, O.A., Shepherd, A.K., Wasserman, S.A., Kim, S.M., and Wang, J.W. (2015). Starvation promotes concerted modulation of appetitive olfactory behavior via parallel neuromodulatory circuits. Elife 4.

[23] Komatsu, H., Mori, I., Rhee, J.S., Akaike, N., and Ohshima, Y. (1996). Mutations in a cyclic nucleotide-gated channel lead to abnormal thermosensation and chemosensation in C. elegans. Neuron 17, 707–718.

[24] Kuhara, A., Ohnishi, N., Shimowada, T., and Mori, I. (2011). Neural coding in a single sensory neuron controlling opposite seeking behaviours in Caenorhabditis elegans. Nature communications 2, 355.

[25] Kuhara, A., Okumura, M., Kimata, T., Tanizawa, Y., Takano, R., Kimura, K.D., Inada, H., Matsumoto, K., and Mori, I. (2008). Temperature sensing by an olfactory neuron in a circuit controlling behavior of C. elegans. Science 320, 803–807.

[26] Kunitomo, H., Sato, H., Iwata, R., Satoh, Y., Ohno, H., Yamada, K., and Iino, Y. (2013). Concentration memory-dependent synaptic plasticity of a taste circuit regulates salt concentration chemotaxis in Caenorhabditis elegans. Nature communications 4, 2210.

[27] Luo, L., Cook, N., Venkatachalam, V., Martinez-Velazquez, L.A., Zhang, X., Calvo, A.C., Hawk, J., MacInnis, B.L., Frank, M., Ng, J.H., et al. (2014). Bidirectional thermotaxis in Caenorhabditis elegans is mediated by distinct sensorimotor strategies driven by the AFD thermosensory neurons. Proceedings of the National Academy of Sciences of the United States of America 111, 2776–2781.

[28] Mayford, M., Siegelbaum, S.A., and Kandel, E.R. (2012). Synapses and memory storage. Cold Spring Harb Perspect Biol 4.

[29] Mohri, A., Kodama, E., Kimura, K.D., Koike, M., Mizuno, T., and Mori, I. (2005). Genetic control of temperature preference in the nematode Caenorhabditis elegans. Genetics 169, 1437–1450.

[30] Mori, I., and Ohshima, Y. (1995). Neural regulation of thermotaxis in Caenorhabditis elegans. Nature 376, 344–348.

[31] Narayan, A., Laurent, G., and Sternberg, P.W. (2011). Transfer characteristics of a thermosensory synapse in Caenorhabditis elegans. Proceedings of the National Academy of Sciences of the United States of America 108, 9667–9672.

[32] Okochi, Y., Kimura, K.D., Ohta, A., and Mori, I. (2005). Diverse regulation of sensory signaling by C. elegans nPKC-epsilon/eta TTX-4. EMBO J 24, 2127–2137.

[33] Rabinowitch, I., Chatzigeorgiou, M., Zhao, B., Treinin, M., and Schafer, W.R. (2014). Rewiring neural circuits by the insertion of ectopic electrical synapses in transgenic C. elegans. Nature communications 5, 4442.

[34] Ramot, D., MacInnis, B.L., and Goodman, M.B. (2008a). Bidirectional temperature-sensing by a single thermosensory neuron in C. elegans. Nature neuroscience 11, 908–915.

[35] Ramot, D., MacInnis, B.L., Lee, H.C., and Goodman, M.B. (2008b). Thermotaxis is a robust mechanism for thermoregulation in Caenorhabditis elegans nematodes. The Journal of neuroscience: the official journal of the Society for Neuroscience 28, 12546–12557.

[36] Root, C.M., Ko, K.I., Jafari, A., and Wang, J.W. (2011). Presynaptic facilitation by neuropeptide signaling mediates odor-driven food search. Cell 145, 133–144.

[37] Son, H., Madelian, V., and Carpenter, D.O. (1996). The translocation and involvement of protein kinase C in mossy fiber-CA3 long-term potentiation in hippocampus of the rat brain. Brain Res 739, 282–292.

[38] Takeishi, A., Yu, Y.V., Hapiak, V.M., Bell, H.W., O’Leary T., and Sengupta, P. (2016). Receptortype Guanylyl Cyclases Confer Thermosensory Responses in C. elegans. Neuron 90, 235–244.

[39] Tsukada, Y., Yamao, M., Naoki, H., Shimowada, T., Ohnishi, N., Kuhara, A., Ishii, S., and Mori, I. (2016). Reconstruction of Spatial Thermal Gradient Encoded in Thermosensory Neuron AFD in Caenorhabditis elegans. The Journal of neuroscience: the official journal of the Society for Neuroscience 36, 2571–2581.

[40] Tsunozaki, M., Chalasani, S.H., and Bargmann, C.I. (2008). A behavioral switch: cGMP and PKC signaling in olfactory neurons reverses odor preference in C. elegans. Neuron 59, 959–971.

[41] Wang, D., O’Halloran, D., and Goodman, M.B. (2013). GCY-8, PDE-2, and NCS-1 are critical elements of the cGMP-dependent thermotransduction cascade in the AFD neurons responsible for C. elegans thermotaxis. J Gen Physiol 142, 437–449.

[42] White, J.G., Southgate, E., Thomson, J.N., and Brenner, S. (1986). The structure of the nervous system of the nematode Caenorhabditis elegans. Philosophical transactions of the Royal Society of London Series B, Biological sciences 314, 1–340.

[43] Yau, K.W., and Nakatani, K. (1985). Light-suppressible, cyclic GMP-sensitive conductance in the plasma membrane of a truncated rod outer segment. Nature 317, 252–255.

[44] Yu, Y.V., Bell, H.W., Glauser, D.A., Van Hooser, S.D., Goodman, M.B., and Sengupta, P. (2014). CaMKI-dependent regulation of sensory gene expression mediates experience-dependent plasticity in the operating range of a thermosensory neuron. Neuron 84, 919–926.

[45] Zisopoulou, S., Asimaki, O., Leondaritis, G., Vasilaki, A., Sakellaridis, N., Pitsikas, N., and Mangoura, D. (2013). PKC-epsilon activation is required for recognition memory in the rat. Behav Brain Res 253, 280–289.

